# Structural and molecular basis of the epistasis effect in enhanced affinity between SARS-CoV-2 KP.3 and ACE2

**DOI:** 10.1101/2024.09.03.610799

**Authors:** Leilei Feng, Zhaoxi Sun, Yuchen Zhang, Fanchong Jian, Sijie Yang, Lingling Yu, Jing Wang, Fei Shao, Xiangxi Wang, Yunlong Cao

## Abstract

The recent emergence of SARS-CoV-2 variants KP.2 and KP.3 has been marked by mutations F456L/R346T and F456L/Q493E, respectively, which significantly impact the virus’s interaction with human ACE2 and its resistance to neutralizing antibodies. KP.3, featuring F456L and Q493E, exhibits a markedly enhanced ACE2 binding affinity compared to KP.2 and the JN.1 variant due to synergistic effects between these mutations. This study elucidated the structures of KP.2 and KP.3 RBD in complex with ACE2 using cryogenic electron microscopy (Cryo-EM) and decipher the structural and thermodynamic implications of these mutations on receptor binding by molecular dynamics (MD) simulations, revealing that F456L mutation facilitates a more favorable binding environment for Q493E, leading to stronger receptor interactions which consequently enhance the potential for incorporating additional evasive mutations. These results underscore the importance of understanding mutational epistatic interactions in predicting SARS-CoV-2 evolution and optimizing vaccine updates. Continued monitoring of such epistatic effects is crucial for anticipating new dominant strains and preparing appropriate public health responses.

## Introduction

Recently, the SARS-CoV-2 KP.2-like (known as “FLiRT”) and KP.3 variants from the JN.1 family have become the dominant variants. KP.2 and KP.3 carries new F456L/R346T and F456L/Q493E mutations on the viral receptor-binding domain (RBD) of the Spike glycoprotein compared to JN.1, respectively (Fig. 1a-b) ^1–8^. Mutations observed in these lineages, especially F456L and Q493E, are located on the interface between RBD and human ACE2 (hACE2), and are shown to affect the receptor-binding affinity and substantially escape the Class 1 neutralizing antibodies (NAbs) ^9–12^.

**Fig. 1.**
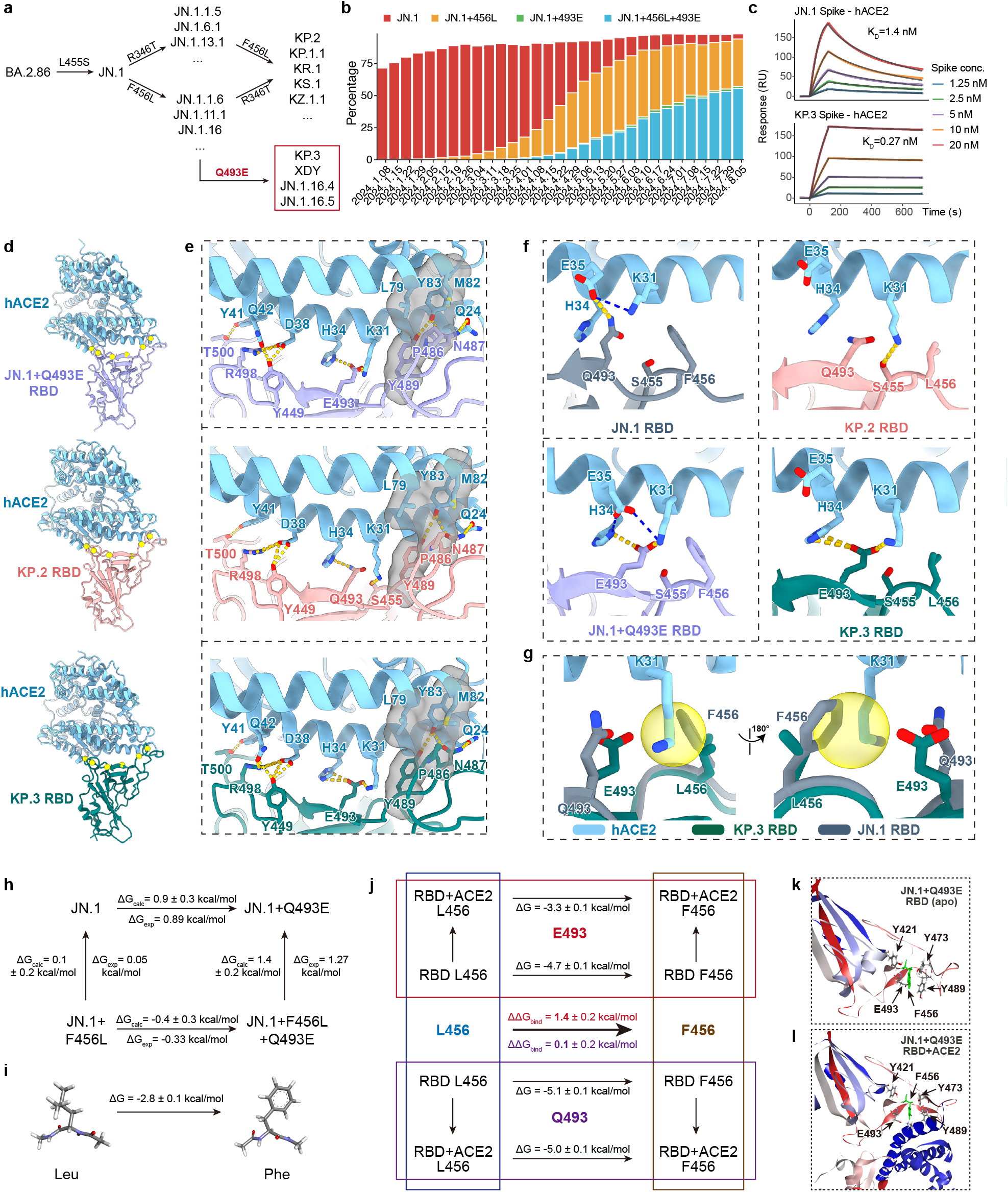
Structural and molecular basis of the enhanced receptor binding in SARS-CoV-2 KP.3. **a**, Convergent emergence of SARS-CoV-2 RBD substitutions located on the receptor-binding interface (R346T, F456L, Q493E) in multiple JN.1-derived sublineages. **b**, Percentage of JN.1 (F456+Q493), JN.1+F456L (Q493), JN.1+Q493E (F456), and JN.1+F456L+Q493E sequences from Jan 2024 to Aug 2024. Data were collected from GISAID. **c**, SPR sensorgrams for the hACE2 binding of JN.1 and KP.3 Spike. Representative results of two replicates are shown. **d**, Overall structures of complexes of JN.1-Q493E RBD, KP.2 RBD, KP.3 RBD in complex with hACE2, purple, pink, green cartoons represent JN.1-Q493E, KP.2, KP.3 RBD respectively, blue color represents hACE2. Residues involved in interactions with hACE2 are shown as yellow balls. **e**, Overall interactions of three variants RBDs with hACE2. Contacting residues are shown sticks, hydrogen bonds are shown as yellow dotted lines. **f**, Comparations of interfaces around site 456 and 493 of JN.1RBD-hACE2 (8Y18), JN.1-Q493E RBD -hACE2, KP.2 RBD-hACE2, KP.3 RBD-hACE2 complexes. Grey cartoons represent JN.1 RBD, other colors indicate as previous in panel **a. g**, Superimposition of hACE2-binding interfaces of JN.1 RBD-hACE2 and KP.3 RBD-hACE2 complexes. Potential clash that limits side chain conformation is shown in yellow circles. **h**, Mutation-induced ACE2-binding affinity variation between different mutants computed with alchemical free energy calculation. **i**, The free energy variation of the L-to-F mutation in capped residues (ACE-X-NME). The computed value -2.8 kcal/mol agree well with the experimental value-3 kcal/mol. **j**, In L-to-F mutations, leg-specific free energy changes along with the net variations of ACE2-binding affinities computed with the cycle-closure condition. The change of RBD-ACE2 binding affinity is estimated indirectly as the difference of free energy variations along the RBD-ACE2 complex leg and the solvated RBD leg, i.e.,*ΔΔG* = *ΔG*_mutation, complex_ −*ΔG*_mutation, RBD_. **k**, The simulation relaxed apo RBD with marks on the central 456 residue and its spatial neighbors contributing to stabilization effects. **l**, The RBD-ACE2 complex with marks highlighting the stabilizing tyrosine residues and the cavity-forming (RBD) E493-K31 (ACE2) coordination.

Notably, the enhanced hACE2 binding and immune evasion of KP.3-like variants compared to JN.1 and KP.2-like variants suggest their global prevalence in the near future ^13^. Interestingly, while the F456L mutation minimally perturbs the ACE2 affinity under Q493 in the case of KP.2, a near 5 to 10-fold increase of ACE2 binding affinity is observed under E493 for KP.3, demonstrating a epistasis effect between L456 and E493 (Fig. 1c, Supplementary Table S1-S2) ^13,14^. The higher receptor-binding affinity achieved by epistasis may lead to higher tolerance to additional evasive mutations, enhancing the viral evolutionary potential of KP.3 ^10,16,17^. However, the structural and molecular insights of this epistasis effect between KP.3 and human ACE2 remain absent. It is crucial to decipher the molecular basis of this L456-E493 mutational cooperativity to help understand the future evolution potential of KP.3-like variants and optimize the update of SARS-CoV-2 vaccines. Here we report the structure of KP.2 and KP.3 RBD in complex with ACE2 along with the intermediate, JN.1+Q493E, using cryogenic electron microscopy (Cryo-EM). Enabled by extensive molecular dynamics (MD) simulations, we estimated the mutation-induced ACE2 affinity variation, probed the protein dynamics, and elucidated the underlying physical chemistry of the observed ACE2-binding cooperativity in the F456L and Q493E mutations.

## Results

To elucidate the impacts of mutations Q493E and F456L on the binding affinity disparities between JN.1, KP.2 and KP.3 RBD with ACE2, particularly the synergistic effects of F456L and Q493E, we constructed recombinant subunit proteins of JN.1+Q493E, KP.2 (JN.1+R346T+F456L) and KP.3 (JN.1+F456L+Q493E) RBD and determined the variants’ structures in complex with hACE2 at a resolution of 3.30 Å, 3.30 Å and 3.29 Å using Cryo-EM (Fig. 1d and Supplementary Fig. S1). The unambiguous high-resolution electron densities allow for reliable reconstruction and analyses of the RBD-hACE2 interaction interfaces. Consistent with previous reports on earlier SARS-CoV-2 variants, the interacting residues of the three RBDs with hACE2 is comparable, including Y449, N487, Y489, R498, T500 (Fig. 1e) ^18^. Precisely, N487 in three RBDs forms a hydrogen bond (H-bond) with Y83 and Q24 of hACE2 respectively, and Q24 of hACE2 forms an H-bond with Y489 of RBDs. In addition, L79, M82, and Y83 of hACE2 as well as P486 and Y489 of all three RBDs construct a hydrophobic pocket, further stabilizing the interaction. Y41 of hACE2 interacts with T500 in three RBDs through an H-bond. However, the interactions surrounding locus 493 varies between three RBDs. In JN.1+Q493E/hACE2 complex, E493 of RBD forms one salt bridge with H34 and K31 of hACE2 correspondingly, and Y449 nearby forms an H-bond with D38 and Q42 of hACE2 respectively, then R498 binds to Q42 of hACE2 through two H-bonds. While in KP.2/hACE2, Q493 of RBD binds to H34 of hACE2 through an H-bond, meanwhile, Y449 and R498 form two H-bonds with D38 separately. Uniquely for KP.2 RBD, an H-bond was introduced between RBD S455 and hACE2 K31. Importantly, KP.3 RBD shows a higher hACE2 affinity compared with KP.2 RBD and JN.1+Q493E RBD. E493 of KP.3 RBD forms two salt bridges with hACE2 H34 and one with K31. Moreover, Y449 binds to hACE2 D38 and Q42 through three H-bonds, and R498 also forms two H-bonds with D38 and Q42, facilitating a more compact contact (Fig. 1f).

It is reported that residue 346 regulates R493 but not Q493 on interaction surface through long-range alterations in the RBD ^19^. We compared the structure of JN.1 RBD(R346, Q493)/hACE2, KP.2 RBD (T346, Q493)/hACE2 and found that Q493 and nearby residues of two RBDs adopt comparable conformation (Supplementary Fig. S2A). In this case, we infer that R346T substitution does not visibly influence interactions, and that KP.2 interacts with RBD similar to JN.1+F456L in the analyses, which is consistent with the experimental affinity measurements reported recently. Meanwhile, alignment of KP.2 RBD/hACE2 (T346, Q493) and KP.3 RBD/hACE2 (R346, E493) shows that Q493 of KP.2 RBD adopts similar orientation to E493 of KP.3 RBD (Supplementary Fig. S2B), which might indicate that E493 is not affected by residue 346 either. Considering that R493 possesses a longer side chain than Q493 and E493, we speculate that the length of side chain of residue 493 is a critical prerequisite for the synergetic effect between residue 346 and 493. In R493 background, when R346 pushes the backbone of RBD on the interface slightly closer to hACE2,the elongated side chain of R493 could insert into the cavity of H34, E35 and D38 of hACE2 and form considerable interactions ^18,19^. While in Q/E493 RBDs, the short side chain of glutamine or glutamic acid is not able to contact with three residues of hACE2 simultaneously, therefore the synergetic effect could not be realized in Q/E493 background.

To further delineate the effects of loci 456 and 493, we anatomized the binding interface with ACE2 around the two residues in JN.1, JN.1+Q493E, KP.2 and KP.3 RBD, respectively. In the JN.1 RBD-ACE2 complex, the strong steric hinderance of F456 “pushes” K31 closer, forming a salt bridge to E35 in ACE2, which caused E35 manifest a rotamer conformation point towards Q493 of RBD (Fig. 1f). Consequently, a hydrogen bond was introduced between RBD Q493 and E35. When F456 is substituted by L456 in KP.2 RBD, it points to the opposite side and does not influence K31 of ACE2, therefore ACE2 K31 inserts between Q493 and L456 and is able to form a hydrogen bond with S455 of RBD. Meanwhile E35 of ACE2 springs back and lose contact with RBD Q493. Therefore, the substitution F456L alone does not incur pronounced alteration of hydrogen bond network RBD and ACE2, which is also consistent with the moderate ACE2-binding affinity change when comparing JN.1 and KP.2 RBD ^13,14^. In JN.1+Q493E RBD, F456 maintains the same conformation as in JN.1, hence E35 forms two salt bridges with H34 and K31 in ACE2, further stabilizing the rotamer orientation which points to RBD E493. As a result, although substitution from glutamine to positively charged glutamic acid enables two salt bridge, the electrostatic repulsion E493 of RBD with ACE2 E35 places E493 in a dilemma, resulting in significantly decreased interaction. For KP.3 RBD harboring an additional F456L, in contrast to JN.1+Q493E, the L456 adopts the same rotamer orientation as KP.2 RBD, allowing space for ACE2 K31 to insert (Fig. 1g). As a result, E493 forms three salt bridges with ACE2 H34 and K31 and avoids electrostatic clash with ACE2 E35 in parallel. The superimposition of JN.1 onto KP.3 RBD shows pronounced steric clashes between K31 of ACE2 and F456 of RBD, thereby disturbing the binding mode. Thus, F456L could be a prerequisite for the fitness of Q493E.

While the analyses of Cryo-EM structures describe the detailed RBD-hACE2 interactions, the underlying thermodynamic driving forces need further investigation, considering the neglection on the apo dynamics of the RBDs and the absence of entropic estimates that play a critical role due to entropy-enthalpy compensation ^20,21^. To provide a faithful thermodynamic picture, we explore the protein dynamics further with MD simulations. Detailed explanations of the simulation procedure are provided in section S2. In each mutational transformation, the alchemical free energy calculation requires the calculation along two transformation pathways (the solvated RBD *ΔG*_mutation, unbound_ and the RBD-ACE2 complex *ΔG*_mutation, bound_), which are then combined to estimate the mutation-induced ACE2-affinity variation according to the cycle-closure condition, i.e., *ΔΔG* = *ΔG*_binding, mutated_ −*ΔG*_binding, original_ = *ΔG*_mutation, bound_ −*ΔG*_mutation, unbound_. Satisfactory convergence behaviors with respect to the equilibration time are observed for both solvated RBD and RBD-ACE2 complex legs under both E493 and Q493 backgrounds (Supplementary Fig. S3).

The thermodynamic impacts of single-site mutations from both alchemical free energy calculations give consistent results compared with the corresponding experimentally measured values (Fig. 1h). Out of the four single-site mutations, the most critical one is the L456F with E493 (JN.1+Q493E vs. JN.1+F456L+Q493E), which exhibits a huge ∼10-fold K_D_ drop in SPR experiments. By contrast, this process with Q493 leads to a close-to-zero affinity variation. These crucial trends are correctly captured by molecular simulations (agrees within uncertainty), thus validating the reliability of the simulation outcome. For the specific mutation of the greatest interest, the L-to-F mutation at the 456 site, we investigate the driving force of the 493-dependent behavior in great details. We perform an additional simulation mutating L-to-F in a capped peptide, i.e., ACE-X-NME with X being Leu or Phe, which gives a free energy variation of -2.8 ± 0.1 kcal/mol (Fig. 1i). This value is in good agreement with the difference between solvation free energies of the side-chain analogues observed experimentally (−3.0 kcal/mol) ^22^. The thermodynamic driving force of this negative (or F-preferred) free energy variation is mainly due to the distinct hydration of the central Leu or Phe residue. We calculated the free energy variations along with the net change of ACE2 affinity caused by L456F substitution within both E493 and Q493 background (Fig. 1j). Compared with the capped residue, all L-to-F transformations are accompanied by more favorable (negative) free energy changes, which suggests the existence of additional stabilizing effects in the protein environment. The additional new interaction components added in the solvated RBD are contributed by intra-molecular interactions. Three critical residues with aromatic rings are in the spatial neighborhood of the 456 site (Fig. 1k). Specifically, tyrosine residues at RBD 421, 473, and 489 sites could stack with the F456 site, stabilizing the L-to-F substitution and shifting the L-to-F free energy variation from the -2.8 kcal/mol value in capped residues to more negative -4.7 kcal/mol (E493) and -5.1 kcal/mol (Q493) in the protein environment. Compared with the unbound solvated RBD state, additional components involved in the bound state are the intermolecular RBD-ACE2 coordination. The formation of such interfacial contacts rearranges both the protein conformation and the solvation environment. For the specific 456 site, the residue lies in the pocket formed by the aromatic residues (tyrosine), the 493 residue on the RBD side as well as several residues on the ACE2 side (Fig. 1l). The critical cavity-forming residue related to the E493 and Q493 is K31 on the ACE2 side. The intermolecular coordination between the 493 site of RBD and K31 of ACE2 is more stable (and thus stronger) with E493 due to the favorable interaction between explicit charges, which elevates the energetic penalty of enlarging the pocket to encapsulate the larger side chain of Phe (Supplementary Video S1 and S2).

Overall, the significant difference between the ACE2 affinity variation upon F456L with Q493 and that with E493 is a joint effect of molecular solvation (L-vs-F solvation tendency), the intra-molecular pi-pi stacking stabilization provided by the neighboring tyrosine residues of RBD, the intermolecular Q493-K31 and E493-K31 coordinations that alter the cavitation cost of enlarging the cavity size from the L456 to the larger F456, in addition to some conformational rearrangements of the secondary structures and the rearrangement RBD-ACE2 backbone packing (Supplementary Fig. S4). Although the F456L substitution alone does not affect hACE2 binding affinity and Q493E diminishes affinity, the combination of F456L and Q493E shows a surprisingly epistasis on RBD-hACE2 interaction. Similar synergistic effects have also been observed in the L455F+F456L “FLip” mutations, where F456L is also a prerequisite of L455F mutation, indicating that F456L is an interesting and necessary immediate during SARS-CoV-2 evolution ^10^. These findings explain the mechanism for the epistatic interactions between F456L and Q493E substitutions regarding ACE2-binding in KP.3 RBD, and underscore that combination of viral mutations on the receptor-binding motif may preserve receptor-binding affinity while attenuating the activity of receptor-mimicking NAbs. Thorough investigation of the potential epistatic interactions of multiple mutations at the RBD-ACE2 binding interface is imperative, and this phenomenon must be vigilantly monitored to facilitate the precise forecasting of SARS-CoV-2 adaptive evolution and the timely alert of emerging prevalent strains.

## Data availability

The structure models of RBD-ACE2 complexes have been deposited in the protein data bank (PDB) under accession codes 9IUU (JN.1+Q493E), 9IUQ (KP.2) and 9IUP (KP.3).

## Acknowledgements

We express our gratitude to Drs. Boling Zhu, Xujing Li, and Lihong Chen for their valuable contribution in Cryo-EM data collection at the Center for Biological Imaging (CBI) of the Institute of Biophysics. This work was supported by the Ministry of Science and Technology of China (2021D0102, CPL-1233 and SRPG22-003), the National Key R&D Program (2023YFC3041500, 2023YFC3043200, 2023YFC2306000 and 2018YFA0900801), CAS (YSBR-010) and the National Science Foundation Grants (12034006, 32325004 and T2394482). Yunlong Cao was supported by the National Science Fund for Excellent Young Scholar (32222030). Xiangxi Wang was supported by the National Science Fund for Distinguished Young Scholar (32325004) and the NSFC Innovative Research Group (81921005).

## Competing interests

Y.C. is a co-founder of Singlomics Biopharmaceutials. Other authors declare no competing interests.

## Author Contributions

Y.C. and X.W. conceived and supervised the study. L.Y., J.W., and F.S. constructed the recombinant proteins. L.F. and Y.Z. performed the Cryo-EM experiments and data analysis. Z.S. performed MD simulations. L.F., Z.S., F.J., and S.Y. performed bioinformatic analysis and illustration. L.F., Z.S., and F.J. wrote the manuscript with input from all authors.

## Supplementary information

The Supplementary Information includes the detailed methods and workflow depositing the crystal structures, the procedure of model construction and simulations in molecular modelling and detailed analyses of structural differences of the four mutants involved in the 456-493 mutational cooperativity.

