## Supplementary material for "Structural and molecular basis of the epistasis effect in enhanced affinity between SARS-CoV-2 KP.3 and ACE2": Table S1-S2, Figure S1-S4

**Supplementary Information**

**Table S1.** SPR measurements of dissociation constants and binding affinities for key mutants to hACE2 taken from reference.^1^

| Mutants | K_D_ (nM) | ΔG (kcal/mol) |
| --- | --- | --- |
| JN.1 | 13 | -10.71 |
| JN.1+F456L | 12 | -10.76 |
| KP.2 (JN.1-F456L+R346T) | 11 | -10.81 |
| JN.1+Q493E | 59 | -9.82 |
| KP.3 (JN.1-F456L+Q493E) | 6.9 | -11.09 |

**Table S2.** JN.1 and KP.3 Spike – hACE2 binding kinetics measured by SPR.

| Protein | k_a_ (1/Ms) | k_d_ (1/s) | K_D_ (nM) |
| --- | --- | --- | --- |
| JN.1 Spike | 2.88×10^6^ | 4.01×10^-3^ | 1.39 |
|  | 3.02×10^6^ | 4.04×10^-3^ | 1.34 |
| KP.3 Spike | 3.64×10^5^ | 8.24×10^-5^ | 0.227 |
|  | 3.48×10^5^ | 1.11×10^-4^ | 0.318 |

**S1. Supplementary Methods**

**S1.1. Protein expression and purification.**

KP.2 RBD (I332V, G339H, R346T, K356T, S371F, S373P, S375F, T376A, R403K, D405N, R408S, K417N, N440K, V445H, G446S, N450D, L452W, L455S, F456L, N460K, S477N, T478K, N481K, Δ483, E484K, F486P, Q498R, N501Y, Y505H), KP.3 RBD (I332V, G339H, K356T, S371F, S373P, S375F, T376A, R403K, D405N, R408S, K417N, N440K, V445H, G446S, N450D, L452W, L455S, F456L, N460K, S477N, T478K, N481K, Δ483, E484K, F486P, Q498R, N501Y, Y505H) and JN.1 RBD (I332V, G339H, K356T, S371F, S373P, S375F, T376A, R403K, D405N, R408S, K417N, N440K, V445H, G446S, N450D, L452W, L455S, N460K, S477N, T478K, N481K, Δ483, E484K, F486P, Q498R, N501Y, Y505H) with its mutation Q493E were realized by overlapping PCR with the full-length S gene (residues 1-1208, GenBank: MN908947) as template. The gene of each RBD was constructed into the vector pcDNA3.1 with HIS tags at the C-terminal. RBD was expressed in HEK293F cells as previously described^2^. Cells were centrifuged at 8000 rpm for 15 min and the supernatant was concentrated and exchanged into buffer (20 mM Tris, 200 mM NaCl, pH 8.0) using tangential flow filtration cassette. RBD was captured by Ni-NTA resin and eluted with 200mM imidazole in buffer and verified by SDS-PAGE. Then RBD was further purified by size exclusion chromatography using a Superdex 200 10/300 column (GE Healthcare) pre-equilibrated with buffer (20 mM Tris, 200 mM NaCl, pH 8.0). The N-terminal peptidase domain of human ACE2 was expressed and purified by essentially the same protocol as used for RBD.

**S1.2. Surface plasmon resonance (SPR)**

SPR assays were conducted on Biacore 8K (Cytiva). Human ACE2-Fc was immobilized onto Protein A sensor chips (Cytiva). Purified SARS-CoV-2 JN.1 and KP.3 Spike (S6P+R683A+R685A) samples prepared in serial dilutions (1.25, 2.5, 5, 10, and 20 nM) were injected on the sensor chips. Response units were recorded by Biacore 8K Evaluation Software 3.0 (Cytiva) at room temperature. Responses were fitted to 1:1 binding model to determine the kinetic constants (k_a_ and k_d_), and binding affinities (dissociation equilibrium constant K_D_) using Biacore 8K Evaluation Software 3.0 (Cytiva).

**S1.3. Production of Fab fragment.**

The Fab fragment of purified antibodies were processed using the Pierce FAB preparation kit (Thermo Scientific) as previously described^3^. Briefly, the samples were first applied to desalination columns to remove the salt. And the flow through were collected after centrifugation and incubated with beads attached with papain to cleave Fab fragments from the whole antibody. After that, the mixture were transferred to Protein A columns which binds the Fc fragments. And the Fab fragments were collected and dialyzed into Phosphate Buffered Saline (PBS).

**S1.4. Cryo-EM sample preparation.**

The cryo-EM samples of each RBD in complex with human ACE2 were mixed in a molar ratio of 1:1.2:1.2 (RBD:ACE2:SN1600). The complex was applied to freshly glow-discharged grids (C-flat 1.2/1.3 Au). Grids were blotted for 6 s at 22 °C in 100% relative humidity and plunge-frozen in liquid ethane automatically by Vitrobot (FEI).

**S1.5. Cryo-EM data collection and processing.**

Single-particle cryo-EM data of KP.2 RBD in complex with ACE2 were collected on a 300kV FEI Titan Krios transmission electron microscope equipped with a K3 Summit direct detector. Movies (32 frames, each 0.2 s, total dose of 60 e /Å^-2^) were recorded with a defocus range between 1.5-2.7 μm. Automated single particle data acquisition was carried out by SerialEM, with a calibrated magnification of 96,000, yielding a final pixel size of 0.83 Å. Single-particle cryo-EM data of KP.3 RBD in complex with ACE2 and JN.1-Q493E RBD in complex with ACE2 were collected on a 300kV FEI Titan Krios transmission electron microscope equipped with a Falcon 4 Summit direct detector. Movies (32 frames, each 0.2 s, total dose of 60 e /Å^-2^) were recorded with a defocus range between 1.5-2.7 μm. Automated single particle data acquisition was carried out by EPU, with a calibrated magnification of 96,000, yielding a final pixel size of 0.808 Å.

All datasets were processed using CryoSPARC. For all datasets, movies were beam induced motion corrected and frames averaged. Then, the defocus value of each micrograph was estimated by patch CTF estimation. Particles were autopicked using Template Picker and extracted for further 2D classification and hetero-refinement. Then, 267901 particles of KP.2 RBD-ACE2 complex, 387538 particles of KP.3 RBD-ACE2 complex and 218777 particles of JN.1(Q493E)-ACE2 complex were used for homo-refinement and non-uniform refinement for the final cryo-EM density. The resolutions were evaluated on the basis of the gold-standard Fourier shell correction (threshold = 0.143) and evaluated by ResMap.

**S1.6. Cryo-EM structure model fitting and refinement.**

Initial coordinates were generated by docking reference structures (PDB:8WRL) into cryo-EM density using UCSF Chimera^4^. Then the structures were adjusted and corrected manually in Coot, according to sequences and density. The models were further improved using real-space refinement by PHENIX.

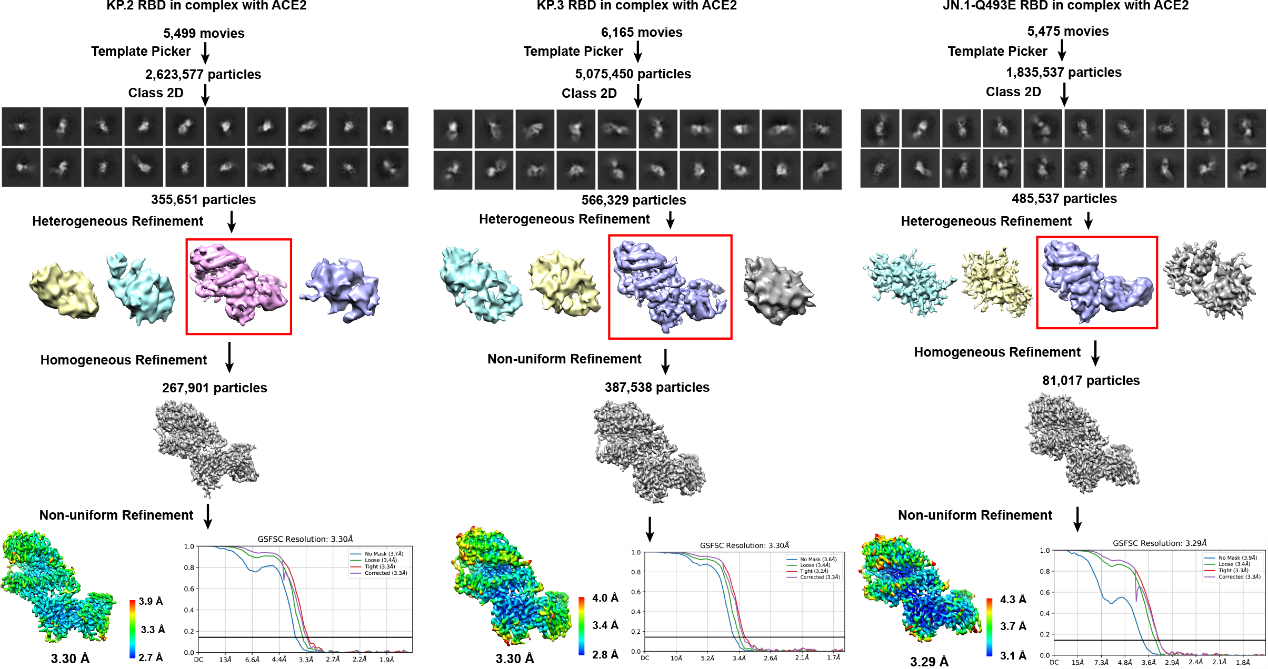

**Fig. S1.** Flowchart illustrating the structure deposition of the RBD-ACE2 complex for KP.2, KP.3 and JN.1-Q493E variants.

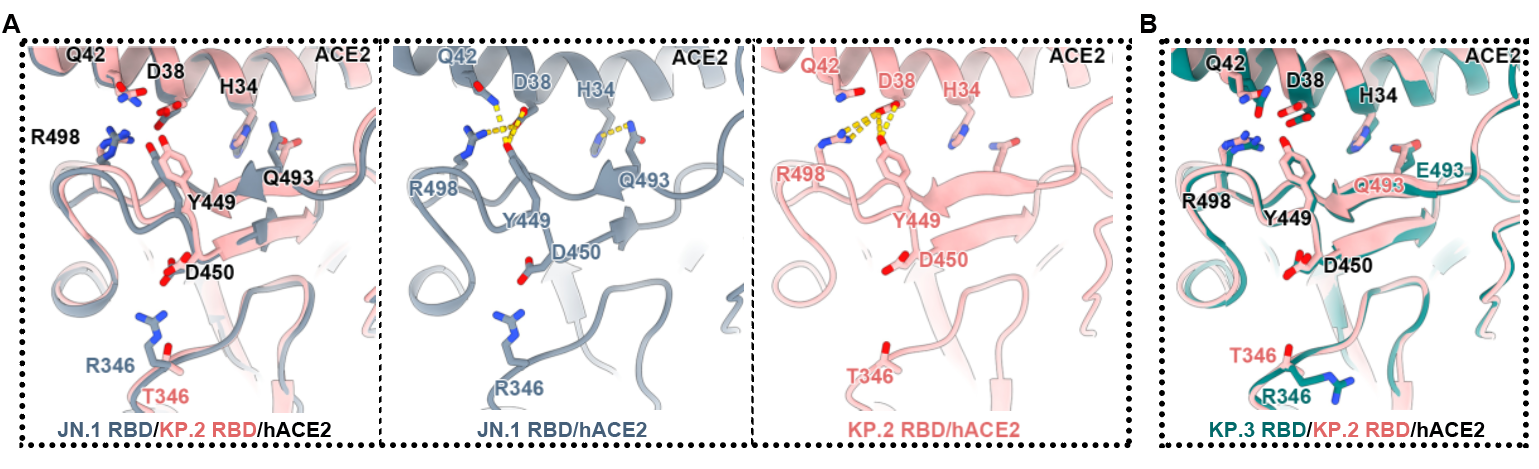

**Fig. S2.** RBD-ACE2 interfacial polar coordinations for JN.1, KP.2, and KP.3 Omicron subvariants.

**S2. Details of molecular simulations.**

Starting from the experimentally deposited structure of the RBD-ACE2 complex of the KP.3 variant, we use homology modelling to construct the 3D structures of the other mutants. We use the three-letter (455-456-493) code for each mutant, and the KP.3, JN.1+Q493E, JN.1 and JN.1+F456L mutants can thus be represented as SL_E, SF_E, SF_Q and SL_Q, respectively. For each system, we first perform pKa predictions with the established PROPKA3 tool^5,6^ to determine the protonation states of ionizable protein residues. We then parametrized the protein system with the AMBER14SB^7^ force field, added disulfide bonds according to the sulfur-sulfur distance of cysteine pairs, neutralized the simulation cell with non-polarizable monovalent spherical counter ions^8,9^ of Na^+^ or Cl^-^, and add TIP3P^10,11^ water with a 12 Å box-edge distance for solvation, creating a simulation box containing more than 140,000 atoms. For each constructed model, we perform 15,000 cycles of geometry minimization, 120 ps constant-volume heating with weak harmonic restraints (3 kcal·mol^-1^·Å^-2^) on heavy atoms of solutes and 2 ns NPT equilibration, after which a 300 ns production run with a sampling interval of 10 ps is initiated to probe the protein dynamics. The 300 ns sampling is separated to 6 juxtaposed 50 ns-length replicates to achieve a better sampling efficiency, following the recommended practice in the recent reference.^12^ The whole simulation procedure is repeated for the unbound state (i.e., solvated RBD).

In the costly alchemical free energy calculations, we consider alchemical transformations between the four variants (i.e., SL_E, SF_E, SL_Q and SF_Q), and in each transformation we consider the variation of a single residue, e.g., L456F or Q493E. The estimation of the mutation-induced ACE2 affinity change requires the simulation of two legs including the solvated RBD (unbound state) and the RBD-ACE2 complex (bound state). The affinity variation is thus obtained indirectly according to the cycle-closure condition, $\Delta\Delta G=\Delta G_{\text{binding, mutated}}-\Delta G_{\text{binding, original}}=\Delta G_{\text{mutation, bound}}-\Delta G_{\text{mutation, unbound}}$. In the non-physical alchemical intermediates, the physically end-state Hamiltonians V_0_ and V_1_ are linearly mixed $V\left( \lambda\right)=\left( 1-\lambda\right)V_{0}+\lambda V_{1}$ , and the soft-core potential with the non-linear separation-shifted scaling regime^13-15^ is applied to avoid vdW singularity. The increment of the alchemical order parameter is set to 0.1 or 0.05. The sampling in each alchemical state includes energy minimization, constant-volume heating, 2 ns NPT equilibration and a final 3*15 ns production with a 4 ps sampling interval, similar to the replicate sampling used in the unbiased simulations of the physical end states. In post-simulation free energy extraction, we first compute the autocorrelation time of the statistical quantify of central interests in alchemical simulations, the partial derivative of the alchemical Hamiltonian ${\frac{\partial U}{\partial\lambda}|}_{\lambda_{i}}$, based on which the time-series data are subsampled with its statistical inefficiency. Then, the statistically optimal multistate free energy estimator MBAR^16^ is employed to obtain the free energy variation between physical end states. A post-simulation finite-size correction^17,18^ is added in situations where the net-charge variation is involved (i.e., Q493E).

In all simulations, we use Langevin dynamics^19^ with the collision frequency of 2 ps^-1^ for temperature regulation at 300 K and the Monte Carlo barostat with isotropic position scaling for pressure regulation at 1 atm. The time step of 2 fs is employed with SHAKE^20,21^ constraints on bonds involving hydrogen. For non-bonded interactions, the PME^22^ method is used to treat long-range electrostatics and the real-space cutoff is set to 10 Å (also the cutoff for vdW interactions). The hybrid-precision (SPFP) GPU engine of pmemd in the AMBER22^23^ suite is used for dynamics propagation.

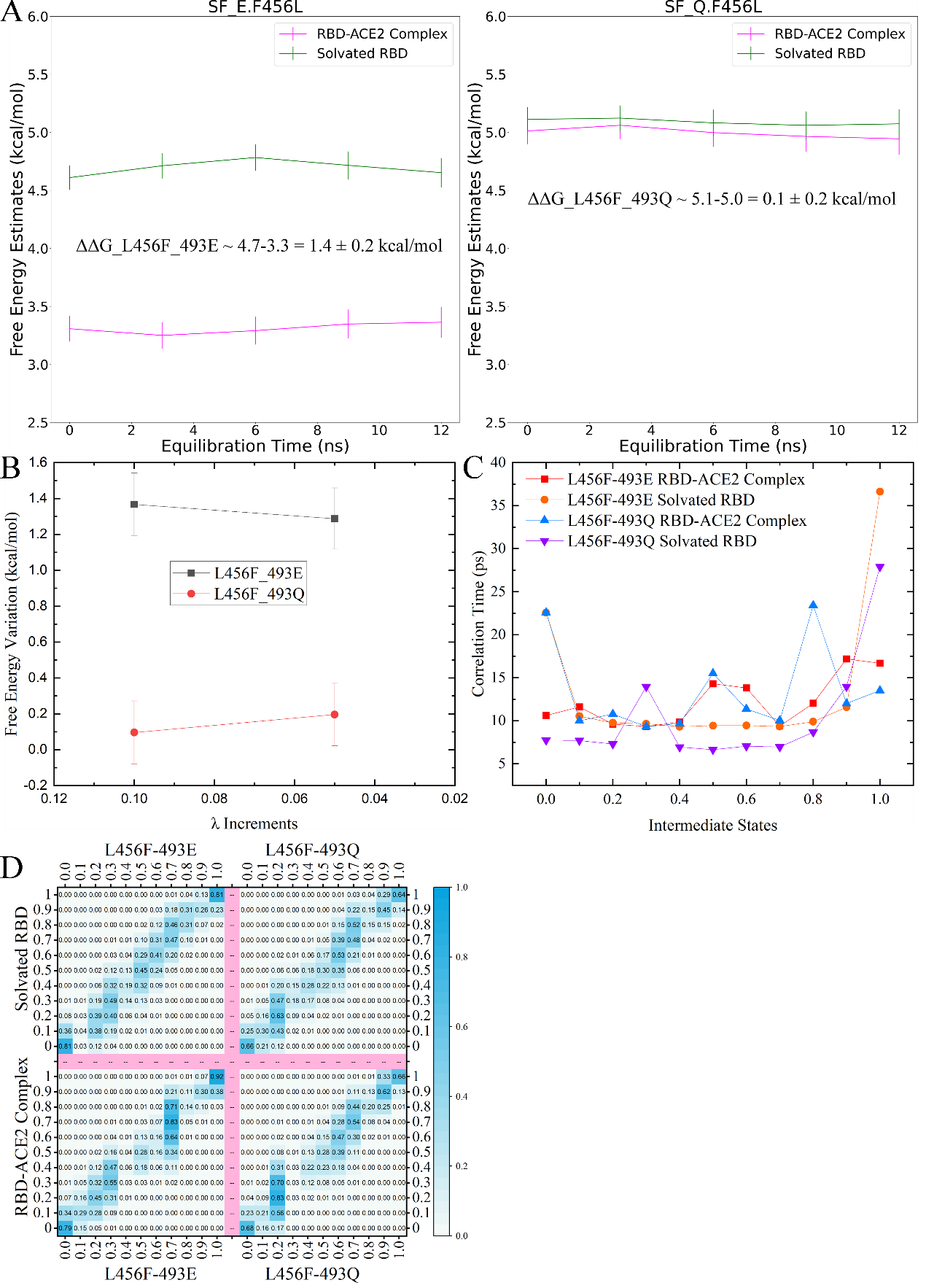

**Fig. S3.** A) Time dependence of free energy changes along the two transformation legs (solvated RBD and RBD-ACE2 complex) during the F456L mutation with 493E and 493Q. The last plateau values are extracted as the free energy variation along each leg. B) L456F mutation-induced affinity variation as a function of the λ increments (or equivalently the number of alchemical intermediate states). C) The correlation time of the partial derivative of the alchemical Hamiltonian ${\frac{\partial U}{\partial\lambda}|}_{\lambda_{i}}$ in all intermediate states. D) The overlap matrix with the 0.1 λ increments quantifying the degree of phase-space overlap between alchemical intermediates.

The time-dependent behaviors of the L456F free energy variations under 493E and 493Q show satisfactory convergence behaviors with respect to the equilibration time are observed for both solvated RBD and RBD-ACE2 complex legs under both E493 and Q493 backgrounds. The λ-increment dependence is shown in panel B, where the 0.1-increment results agree well with the 0.05 increments’ outcome, suggesting the statistical reliability of the computational results. We present the autocorrelation times of the ${\frac{\partial U}{\partial\lambda}|}_{\lambda_{i}}$ in all intermediate states that are used to subsample the time-series data (distilling uncorrelation samples) and the overlap matrix that provides quantitative estimates the phase space overlap, respectively ^18,19^. The main diagonal and its neighbors of the matrix are larger than the empirical 0.03 threshold, validating the reliability of the perturbation-based estimates ^20^.

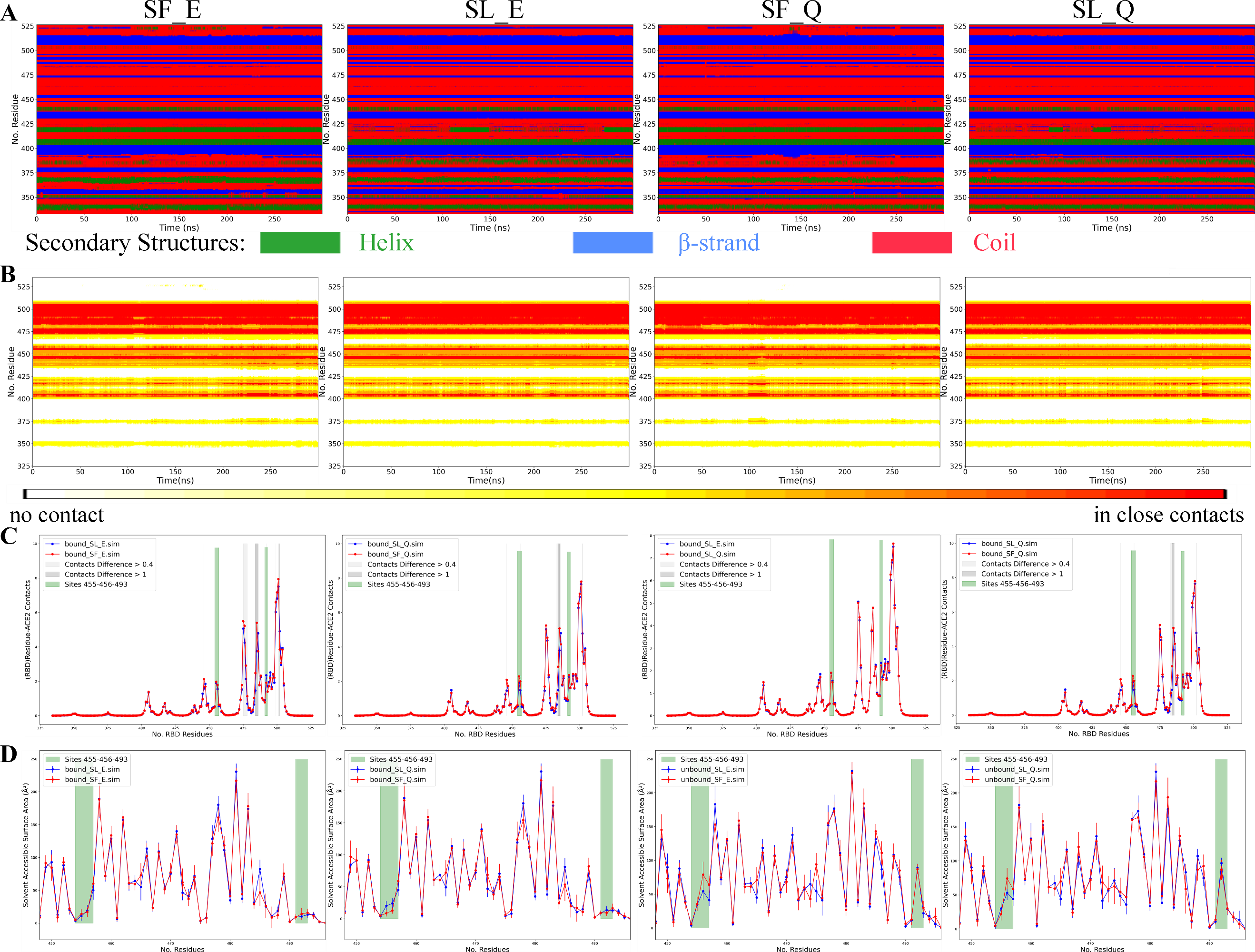

**Fig. S4.** A) Time series of the bound-state secondary structures of RBD with the simplified color code (helix-green, blue-strand and red-coil), B) those of the by-RBD-residue Cα-Cα contacts, C) face-to-face comparisons between the ensemble averaged RBD-ACE2 coordinations, D) the solvent accessible surface area of the key RBD residues in the interfacial contacts in the bound and unbound states. The backbone contacts are computed with the switching function that transits smoothly between 0 and 1, $C_{segment A-segment B}=\sum_{i\in A} \sum_{j\in\text{B}} \frac{1-\left( \frac{r_{ij}}{r_{0}} \right)^{6}}{1-\left( \frac{r_{ij}}{r_{0}} \right)^{12}}$ , where A and B are two groups of atoms and the distance constant r_0_ is set to 8 Å. The solvent accessible surface area is computed using the Shrake-Rupley regime with the golden section spiral sphere and a 1.4 Å probe.

Aside from the residues in direct contacts with the 456 residue, the mutation process is also accompanied with noticeable rearrangements of the other regions of the protein system. According to the time series of the secondary structures of RBD in the bound state shown in Fig. S4A, the L456F mutation leads to the helix formation in the neighborhood of the 383 and the 420 residues and the destabilization of the β-strands around the 486 and 523 residues, regardless of the residue type at the 493 site. Interestingly, while the L456F mutation reshapes the protein conformations, the mutation at the 493 site does not incur obvious conformational rearrangements. The intermolecular contacts between Cα pairs are presented in a by-RBD-residue manner in Fig. S4B, where the backbone coordination also exhibits variations in, e.g., the neighborhood of the 486 residue, upon the L456F mutation. The time-averaged backbone contacts of the four mutants are compared in Fig. S4C. Again, while the L456F mutation leads to variations of intermolecular packing around residues 475-478, 482-486 and 492-502 under both 493E and 493Q, the mutation at the 493 site has little impacts on the intermolecular RBD-ACE2 backbone coordination. In Fig. S4D, we present the face-to-face comparison between solvent-accessible surface area of the key RBD residues involved in the interfacial RBD-ACE2 coordination. In the bound state, the solvent exposures of the four mutants exhibit minor differences, hinting on minor rearrangements of the hydration environment upon mutations. More pronounced changes are observed in the unbound state, where the ACE2 unbinding makes the key residues involved in intermolecular coordinations (e.g., 455-456 and 493) fully exposed to solvent. The significant difference between F456 and L456 is highly related to the size of these molecules and their energetic preference of molecular solvation/hydration.
