## Supplementary figures and images for "Structural and molecular basis of the epistasis effect in enhanced affinity between SARS-CoV-2 KP.3 and ACE2"

### Supplementary Video S1

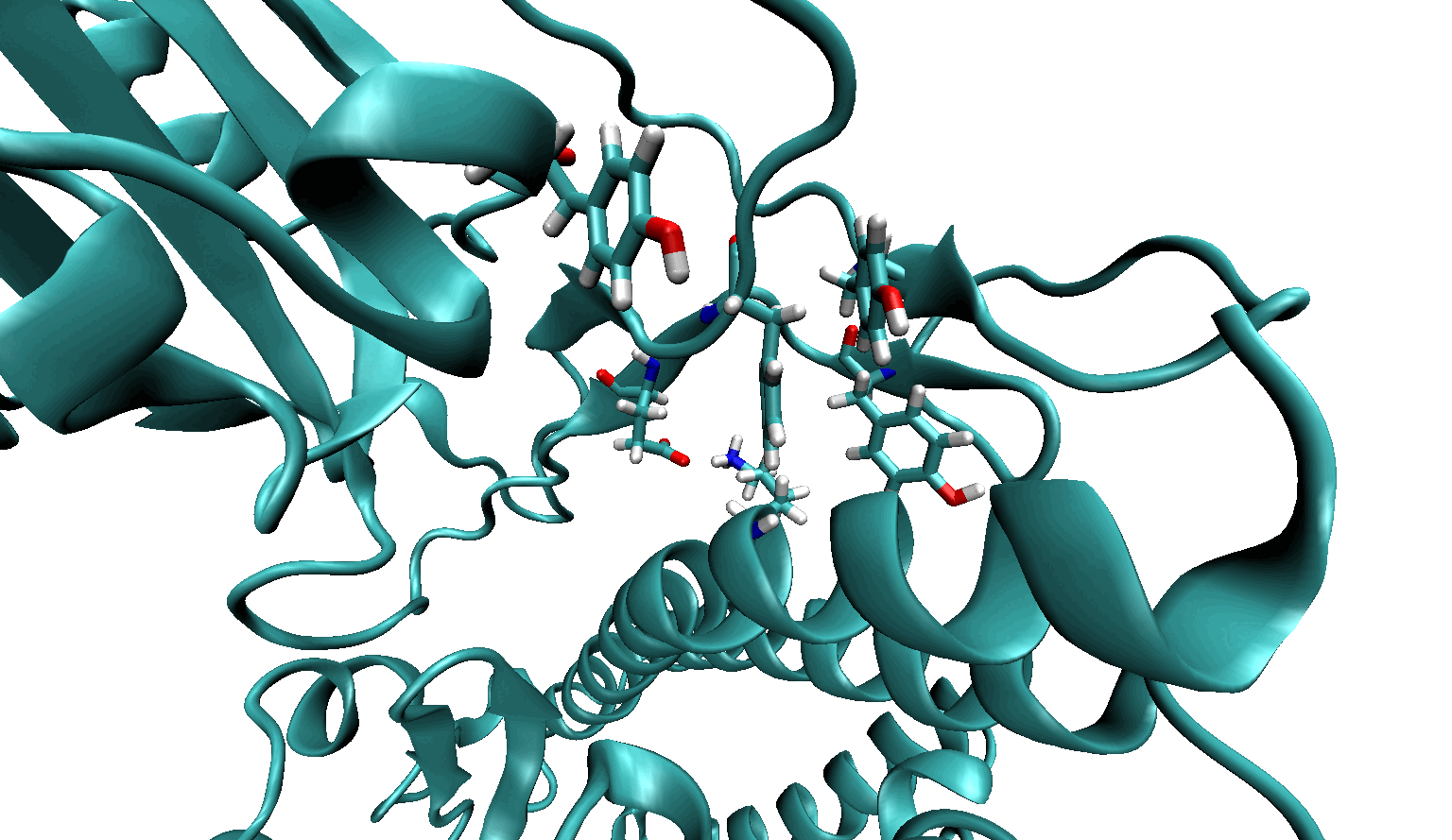

### Supplementary Video S2

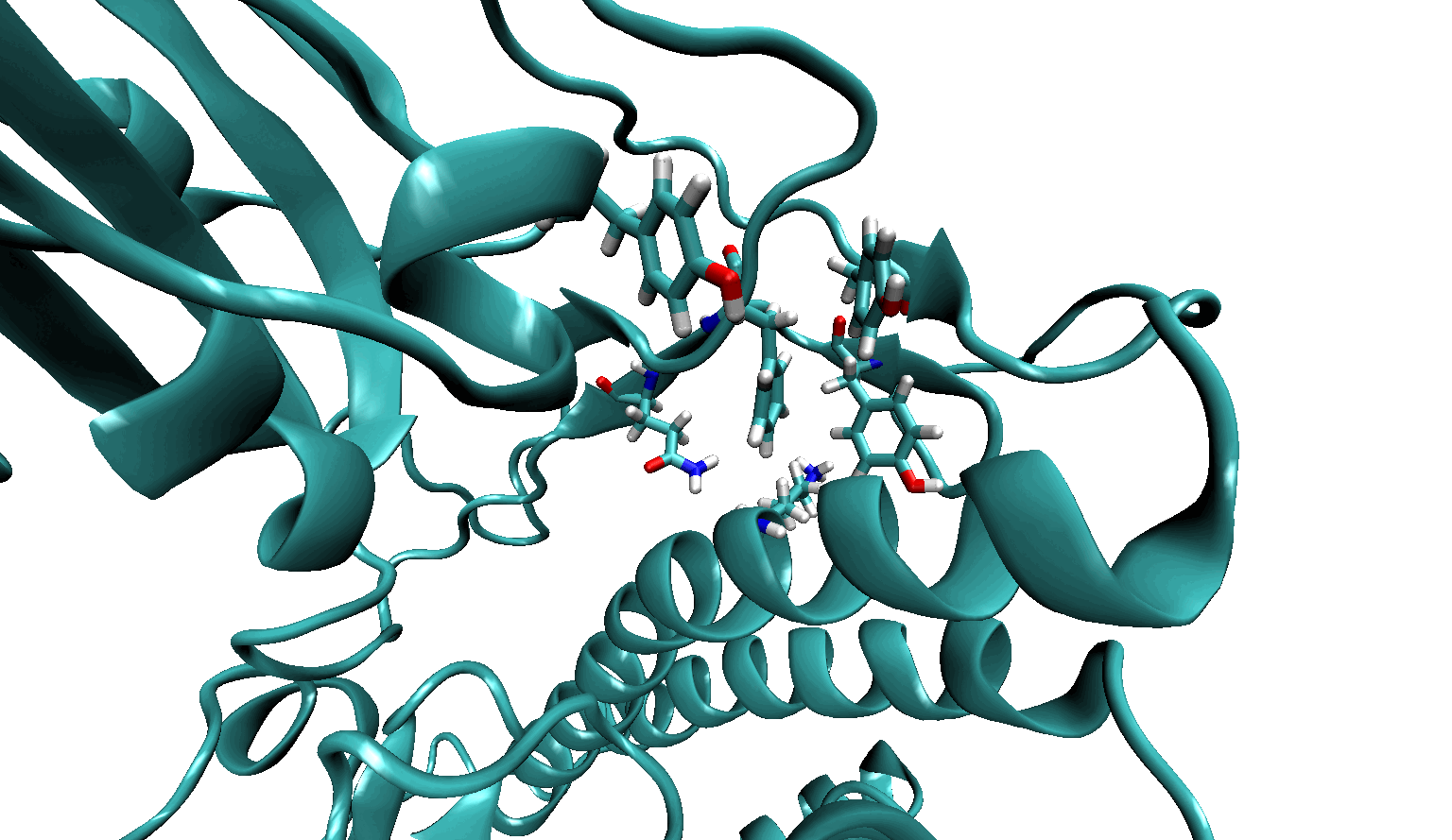
